# SalienceNet: an unsupervised Image-to-Image translation method for nuclei saliency enhancement in microscopy images

**DOI:** 10.1101/2022.10.27.514030

**Authors:** Bouilhol Emmanuel, Edgar Lefevre, Thierno Barry, Florian Levet, Anne Beghin, Virgile Viasnoff, Xareni Galindo, Rémi Galland, Jean-Baptiste Sibarita, Macha Nikolski

## Abstract

Automatic segmentation of nuclei in low-light microscopy images remains a difficult task, especially for high-throughput experiments where need for automation is strong. Low saliency of nuclei with respect to the background, variability of their intensity together with low signal-to-noise ratio in these images constitute a major challenge for mainstream algorithms of nuclei segmentation. In this work we introduce SalienceNet, an unsupervised deep learning-based method that uses the style transfer properties of cycleGAN to transform low saliency images into high saliency images, thus enabling accurate segmentation by downstream analysis methods, and that without need for any parameter tuning. We have acquired a novel dataset of organoid images with soSPIM, a microscopy technique that enables the acquisition of images in low-light conditions. Our experiments show that SalienceNet increased the saliency of these images up to the desired level. Moreover, we evaluated the impact of SalienceNet on segmentation for both Otsu thresholding and StarDist and have shown that enhancing nuclei with SalienceNet improved segmentation results using Otsu thresholding by 30% and using StarDist by 26% in terms of IOU when compared to segmentation of non-enhanced images. Together these results show that SalienceNet can be used as a common preprocessing step to automate nuclei segmentation pipelines for low-light microscopy images.

## 1 INTRODUCTION

Segmentation of cell nuclei is of particular interest for a number of applications such as cell detection, counting or tracking, morphology analysis and quantification of molecular expression. Being able to automatically segment cell nuclei with high precision is particularly important in the case of high-throughput microscopy imaging, where it is often the first step for downstream quantitative data analysis workflows. Indeed, the quality of downstream quantitative analyses is heavily dependent on the accuracy of segmentation, making precise nuclei segmentation essential for drawing meaningful biological conclusions.

Many solutions have been developed, as exemplified by computational competitions such as reported in (Caicedo et al., 2019). Among popular classical image analysis methods used for nuclei segmentation are thresholding and watershed algorithm (Malpica et al., 1997) as well as active contour (Li et al., 2007).

Challenges for automatizing this process are due to a number of image characteristics that can strongly vary between biological and image acquisition conditions. Among them, we can cite such aspects as morphological differences between nuclei from different tissues, heterogeneity of intensity and texture, variation in spatial organization such as the presence of both sparse or dense images with touching nuclei, as well as imaging artifacts (e.g., low signal-to-noise ratio or out-of-focus signal) (Zhou et al., 2019). This results in the necessity to fine-tune numerous parameters between different image acquisitions, or even between individual images.

Recent deep-learning based tools such as Cellpose (Stringer et al., 2021) and StardDist (Schmidt et al., 2018) have greatly reduced the necessity to choose specific parameters. However, despite these important methodological advances, no single combination of methods and parameters can be adopted to automatically perform nuclei segmentation in all images, due to the aforementioned heterogeneity of biological samples and technical artifacts (Hollandi et al., 2022). In particular, live-cell imaging represents a stumbling block for these techniques, since these images are often acquired with low light levels and thus have very low SNR and artifacts. Moreover, current supervised Deep Learning models for nuclei segmentation follow the supervised paradigm and thus require well annotated datasets, which (i) engenders bias due to inaccuracy and incompleteness of available segmentation, where nuclei are improperly annotated and unevenly distributed across images (He et al., 2020) and (ii) limits application to datasets with different image characteristics.

In this paper, instead of focusing on the segmentation itself, we propose to tackle this problem by enhancing the nuclei prior to segmentation step, making the task easy for classical nuclei segmentation tools. Specifically, we take advantage of recent advances in the field of unsupervised generative adversarial networks, aiming to translate images from the source domain to the target domain and alleviating the image annotation requirement. For nuclei enhancement task, the target domain corresponds to images with highly salient nuclei, where strong signal difference between the nuclei and background make segmentation straightforward.

In this work, we introduce SalienceNet, a novel unsupervised Deep Learning-based approach for nuclei saliency enhancement in microscopy images that does not require image annotation when there is need to train the network on new data with different characteristics. We showcase how this can be achieved for translating organoid images acquired with low light into contrasted output images, by training the SalienceNet without providing the network with prior annotation of newly acquired low contrast images. SalienceNet gives a new twist to automatic nuclei segmentation by adapting the domain style transfer framework to this specific task, and thus does not require extensive annotation. We trained a ResNet-based CycleGAN with a custom loss function dedicated to the task of nuclei enhancement, where the intensity of the nuclei and in particular their borders, are made more salient regardless of the contrast, intensity, textures, or shapes of the nuclei in the input data. Furthermore, we evaluated the impact of the obtained nuclei enhancement on the downstream nuclei segmentation by performing segmentation using conventional methods on the enhanced images and have shown that such a pipeline achieves better performance than segmenting the nuclei directly on the original images. We demonstrate here that incorporating SalienceNet in a standard segmentation pipeline, makes it possible to avoid the manual parameter fine-tuning steps.

## 2 RELATED WORK

### 2.1 Nuclei segmentation

Nucleus segmentation methods can be partitioned in two major groups: those that rely on classical image processing approaches and those that propose Deep Learning models. For a thorough review, we refer the reader to (Hollandi et al., 2022).

Image processing pipelines usually contain a number of filtering and thresholding steps combined, if needed, with basic morphological operators to differentiate nuclei (Malpica et al., 1997; Li et al., 2007). A number of such methods are available as plugins of the main biological analyses open-source software tools such as Fiji (Schindelin et al., 2012), ICY (De Chaumont et al., 2012), QuPath (Bankhead et al., 2017) or CellProfiler (McQuin et al., 2018). The development of classical image processing methods for nuclei segmentation is still an active field. For example, in a recent publication (Haase et al., 2020) the authors propose first to denoise the images with Gaussian blur, second to separate regions using Voronöi tessellation, and to finally obtain a binary mask by applying an Otsu thresholding to obtain the segmentation. However, time-consuming parameter finetuning is required from the user at different steps of such classical image processing pipelines, making processing large amount of data impractical (Hollandi et al., 2022).

The need for an automatized solution capable to segment the nuclei in images with different characteristics, pushed for the adoption of methods based on Deep Learning. The U-Net architecture (Ronneberger et al., 2015) is used as part of recent Deep Learning nucleus/cell segmentation methods, such as Cellpose (Stringer et al., 2021) and StarDist (Schmidt et al., 2018). Another successful architecture is Mask R-CNN, that has been recently adapted for nuclei segmentation by the authors of nucleAIzer (Hollandi et al., 2020). ImageJ has recently proposed Deep Learning-based segmentation plugins, and pre-trained models are available through DeepImageJ (Gómez-de Mariscal et al., 2021).

The success of the aforementioned Deep Learning methods for nuclei segmentation is particularly due to the use of large and relatively varied training datasets, with images acquired using different microscopy modalities. Nevertheless, establishing a general solution is still an unmet need, especially for images acquired with novel microscopy techniques, such as for example the live-cell imaging (Ettinger and Wittmann, 2014) that reduces the intensity of the light sources of the microscope to a minimum in order to limit the photo-damage to the cells thus be able to observe them over long periods of time. Resulting images have a reduced signal intensity and low SNR. Importantly, having both (i) not been part of the training and evaluation datasets of the aforementioned methods and (ii) having different characteristics, such images represent a yet unsolved challenge for nuclei segmentation.

Moreover, the performance of the existing supervised deep-learning methods depends on the amount of high-quality annotated data available for training. Despite the large effort that was put to produce publicly available labels for nuclei segmentation, such as the 2018 Data Science Bowl competition, such data is often partially or even incorrectly labeled (He et al., 2020).

### 2.2 Image preprocessing

A frequently used approach to overcome the difficulty of segmentation is to preprocess the images to improve their quality. In the case of nuclei, such enhancement mainly concerns the contrast between the nuclei and the background. Most traditional image enhancement techniques rely on filtering (low pass, high pass) or on naive noise removal such as Gaussian blur. Other methods are based on normalization of image intensity, such as e.g., histogram equalization or contrast stretching (see for review (Qi et al., 2021)). However, in the same way as the segmentation methods themselves, these image preprocessing techniques lack the generalization ability. For example, filtering or signal normalization is not applicable to images with low SNR, as it cannot distinguish well enough the signal from the background.

Deep Learning has been also applied at the preprocessing step, in particular to estimate the transformation function between sets of acquired images and their enhanced counterparts through supervised learning. One of the first and most successful methods was introduced with the CARE network (Weigert et al., 2018), designed to restore fluorescence microscopy data without the need to generate manual training data. The authors showed that it is possible to learn the mapping between low-intensity and high-intensity image pairs using a U-Net based neural network. In the case of live-cell imaging, this makes possible to restore the image quality. However, two characteristics of this network limit the generalization capacity of CARE to new types of images. First, CARE network follows the supervised training paradigm and thus requires matching pairs of the same image and the corresponding nuclei masks, which is time-consuming. Second, CARE comports 5 separately trained networks and uses a disagreement score between the individual network predictions to eliminate unreliable results, which implies that images with characteristics that strongly differ from those in the training set will not be well restored (Weigert et al., 2018).

### 2.3 Image to Image translation

Image quality enhancement has also been approached through image to image translation deep learning methods. The goal is to transform an image having a particular style (source style) into a desired target style. The most efficient models are based on GANs (Pang et al., 2021; Wang et al., 2020). Authors of pix2pix (Isola et al., 2017) were the first to apply a GAN-based architecture to perform the image to image translation. It is a fully supervised method that requires large paired image datasets to train the translation model that transforms the source images to the desired target images. In the context of nuclei segmentation, pix2pix has been used by the authors of nucleAIzer for data augmentation of their training nuclei datasets. Specialized image enhancement models have been since proposed, such as Cycle-CBAM (You et al., 2019) for retinal image enhancement and UW-CycleGAN (Du et al., 2021) for underwater image enhancement, both based on the CycleGAN architecture. Moreover, enhancement of objects of interest has been proposed by the authors of DE-CycleGAN (Gao et al., 2021) to enhance the weak targets for the purpose of accurate vehicle detection.

## 3 PROPOSED METHOD

In this section, we present the SalienceNet nuclei saliency enhancement network in detail. We first overview the network’s architecture, and then we discuss the custom generator loss function that drives the saliency enhancement.

### 3.1 Network architecture

SalienceNet implements the image style transfer for nuclei microscopy images with CycleGAN architecture (Zhu et al., 2017) where the network is composed of two Generative Adversarial Network (GAN) blocks that exchange information during training as shown in figure 1.

**Figure 1:**
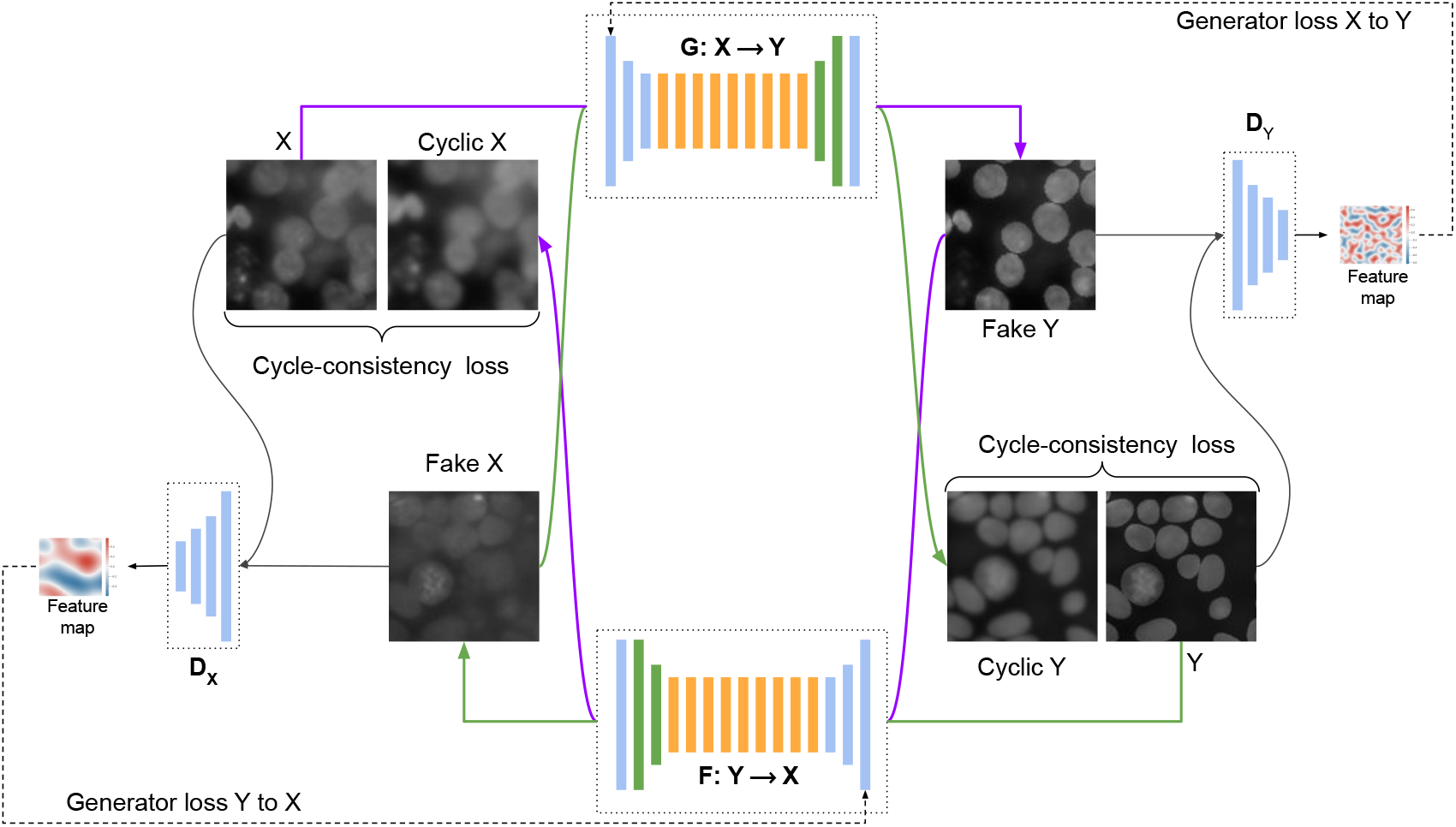
Architecture of SalienceNet. The network is composed by 2 GANs, each GAN having a Generator *G* and a Discriminator *D*. The main element of the Generator is a residual network, while the discriminator is a PatchGAN whose output is a feature map. The generator loss is computed based in this feature map. Cycle consistency loss is computed between the original image *X* and it’s reconstructed image *F*(*G*(*X*)) and between *Y* and it’s reconstructed image *G*(*F*(*Y*)).

Let *X* be the domain of acquired nuclei images and *Y* the style domain of images with enhanced nuclei saliency. Images do not have to be paired. SalienceNet translates an image from domain *X* to the target domain *Y* by learning a mapping *G* : *X* → *Y* such that the distribution of images from *G*(*X*) is indistinguishable from *Y* by an adversarial loss.

The architecture is based on the simultaneous training of two generator models and two discriminator models (see figure 1). First, generator *G* takes input from the domain *X* and outputs images for the target style domain *Y*, and second, generator *F* takes input from the domain *Y* and generates images for the domain *X*. Adversarial discriminator models are used to drive the training by estimating how well the generated images fit the domain: *D*_*Y*_ distinguishes the outputs of *G*(*X*) from domain *Y*; in the same manner, *D*_*X*_ distinguishes the outputs of *F*(*Y*) from domain *X*.

In our model the discriminators *D*_*X*_ and *D*_*Y*_ are implemented as PatchGAN classifiers, composed of 4 convolution blocks (see figure 2), each containing a convolution layer, an instance normalization layer and an activation layer (LeakyReLU). The generators *G* and *F* are implemented as ResNets having the same structure with 3 down convolutions, followed by 9 residual blocks, before applying 2 transpose convolutions and one last convolution layer with a Tanh activation (see figure 2).

**Figure 2:**
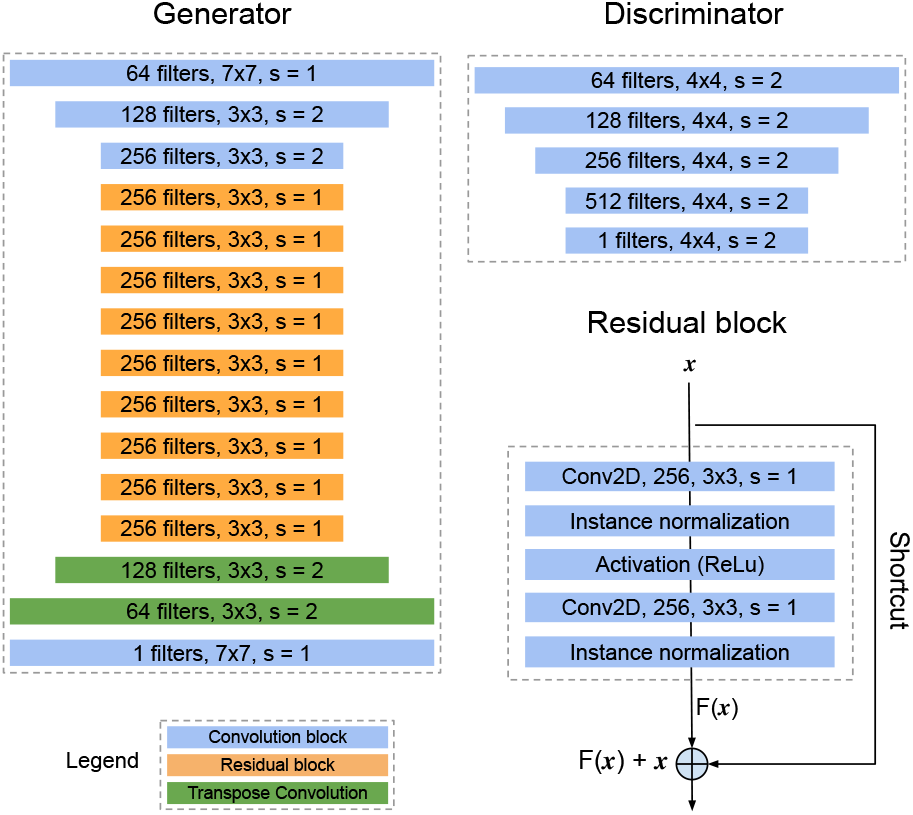
Composition of the networks constituting SalienceNet. Generators embed a residual network composed of 9 residual blocks, each block being itself composed of 2 convolution layers. Convolution blocks are composed of a convolution layer, an instance normalization layer and an activation layer which is ReLU for the generator and LeakyReLU for the discriminator.

The discriminator is implemented as a PatchGAN model that outputs a square feature map of values, each value encoding the probability that the corresponding patch in the input image is real. These values are further averaged to generate the global likelihood.

Since the mapping *G* : *X* → *Y* is highly under-constrained, CycleGAN couples it with an inverse mapping *F*: *Y* → *X* : the output ”fake *Y* ” from the *X* → *Y* generator is used as input to the *Y* → *X* generator, whose output ”cycle *X* ” should match the original input image *X* (and vice versa). This is enforced though the cycle consistency loss to obtain *F*(*G*(*X*)) ≈ *X* and *F*(*G*(*Y*)) ≈ *Y*.

### 3.2 Generator loss function

An additional generator loss is used to enforce the cycle consistency and to measure the difference between the generated output ”cycle *X* ” and *X* as well as between the ”cycle *Y* ” and *Y*. This regularization makes possible to constrain the generation process to image translation.

For SalienceNet we defined the generator loss function as a combination of three terms: (i) the Mean Squared Error (MSE), (ii) the Mean Gradient Error (MGE) and (iii) the Mean Structural SIMilarity index (MSSIM).

The Mean Squared Error (MSE) computes the mean of the squared differences between true and predicted values 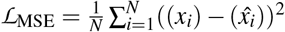. This term ensures that the generator does not produce outliers too far from the target domain. However, MSE used alone is known to lead to blurring due to the averaging between possible outputs, which in image-to-image translation can lead to low-quality blurred results.

In the case of nuclei segmentation, blurring can yield images where nuclei boundaries are particularly difficult to accurately segment. To solve this gradient problem, we added the Mean Gradient Error (Lu and Chen, 2022) term ℒ_MGE_ that measures the differences in edges of objects between two images, with the aim to learn sharp edges. It is based on vertical and horizontal Sobel operators (Kanopoulos et al., 1988), *G*_*v*_ and *G*_*h*_:

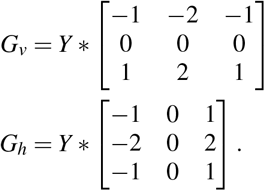

where ∗ is the convolution operator.

These gradients are combined to define a global pixel-wise gradient map 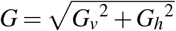. The gradient map for predicted images *Ĝ* is computed in the same way. The ℒ_MGE_ is the defined as:

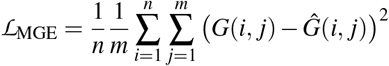

Finally, to drive the network to produce images with a structure similar to the input structure, we added ℒ_MSSIM_ the Mean Structural SIMilarity index (MSSIM) (Wang et al., 2004) as the last term. This loss function compares two images based on luminance *l*, contrast *c* and structural information *s*:

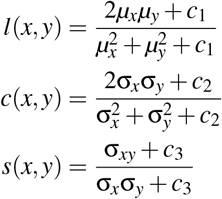

where *µ*_*x*_ and *µ*_*y*_ denote the mean intensity for the input and generated image respectively; σ_*x*_ and σ_*y*_ are standard deviation for the original and generated images; *c*_1_, *c*_2_ and *c*_3_ are constants used to avoid instability when the denominators are close to 0.

The mean SSIM can be obtained over the entire image using a local window as follows:

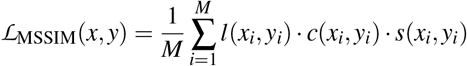

where *x* and *y* denote the input and the generated image, respectively, while *x*_*i*_ and *x* _*j*_ are the images at the *i*-th window when the local window slides over the original and generated images, and *M* is the number of local windows in the image.

In the case of SalienceNet the intuition for the MSSIM loss for the *X* → *Y* generator is to enforce the luminance enhancement, while for the *Y* → *X* generator to preserve the structure.

The total loss function is defined as the weighted sum of the three terms:

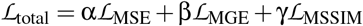

where α, β and γ are the weights of each corresponding term so that the sum α + β + γ = 1.

## 4 DATASETS

To train and evaluate our SalienceNet enhancement method, we have collected different datasets (see figure 3). First, two experimentally acquired and expertly segmented datasets (see section 4.2), which have been previously extensively used for training segmentation models. Second, we have acquired a dataset of organoid images with low-light conditions that specifically represents the segmentation challenge that we want to address, as well as generated the corresponding synthetic datasets (see sections 4.1 and 4.2). These images belong to one of the two styles:

**Figure 3:**
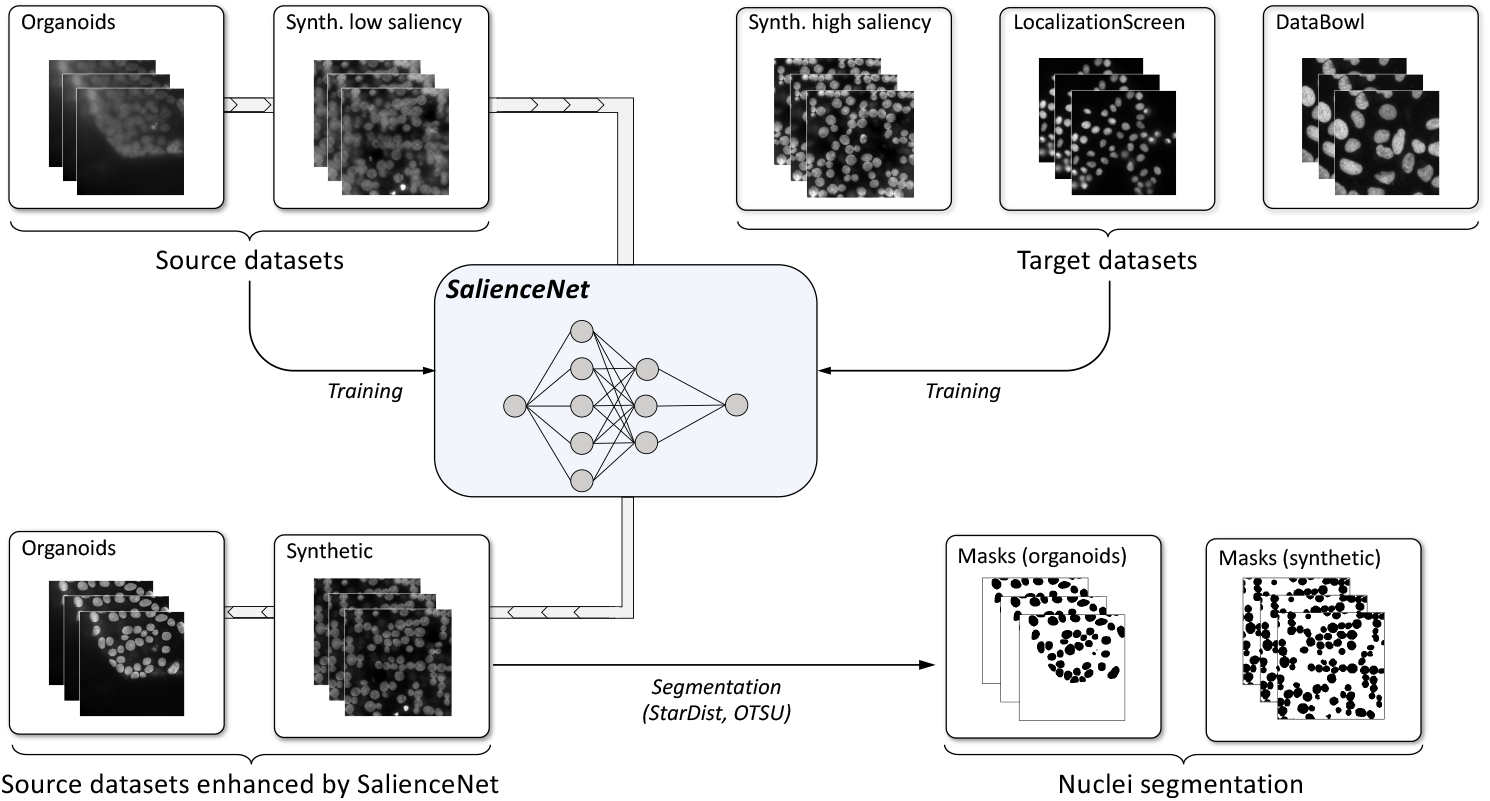
Training and evaluation of SaliencyNet. 90% of source style datasets and of the experimental target style datasets were used to train the network. It was then applied to the remaining 10% of the source style datasets to obtain enhanced images. Nuclei in the images enhanced by SalienceNet were segmented using classical methods with fixed parameters.

1. **Source style** with low saliency of nuclei (organoid and synthetic low saliency datasets),
2. **Target style** with high saliency of nuclei (two experimental datasets and the synthetic high saliency dataset).

### 4.1 Source style datasets

To evaluate whether SalienceNet enables precise nuclei segmentation, we acquired a 3D cell culture dataset with the soSPIM technique. soSPIM is a single objective light-sheet microscopy approach capable of streamlining 3D cell cultures with fast 3D live-imaging at speeds up to 300 3D cultures per hour (Galland et al., 2015; Beghin et al., 2022). We selected 11 neuroectoderm organoids exhibiting a wide variety of shapes and densities. These organoids have been differentiated from hESCs, fixed at day 8, immunostained with DAPI and imaged using soSPIM, yielding 1056 2D slices. These 2D image slices are composing the DS_org_ dataset.

To augment the source style dataset, in addition to the experimental 1056 2D image slices, we generated synthetic images. First, we performed an expert segmentation of nuclei on each individual 2D slice. Second, images paired with their annotated masks were used to train a simple CycleGAN. Finally, this CycleGAN model was applied to transform randomly placed elliptical shapes (roughly approximating nuclei shapes) into organoid look-alike images. The elliptical shapes provide “nuclei” masks in a trivial way. We generated 1500 synthetic low-saliency images, denoted by DS_synth_.

### 4.2 Target style datasets

The goal of our network is to learn to transform an image *i* into *e*(*i*), where saliency (Kim and Varshney, 2006) at nuclei location is enhanced. To provide the target style dataset for training the SalienceNet network, we have collected two experimental datasets where the nuclei saliency was already satisfactory for segmentation by classical pipelines and for which the nuclei segmentation masks are available. We complemented them by a synthetic high-saliency dataset.

In 2018, a Data Science Bowl competition organized by Kaggle released a dataset for a challenge of ”Identification and Segmentation of Nuclei in Cells” of images acquired under different conditions and of different cell types and that vary in size, magnification, and imaging method (brightfield and fluorescence). Nuclei masks have been manually created by specialists and are provided with the dataset. For the purpose of this paper, only grayscale cell culture images were kept, yielding the TS_DB_ with 551 images.

Experimentally acquired nuclei images from human cell lines (Chouaib et al., 2020) were used to define the TS_LS_ dataset. It is composed of 568 images from 57 different acquisition conditions of 32 gene expression measured in the study for the purpose of performing a localization screen. Nuclei masks have been acquired with NucleAIzer.

The synthetic high-saliency dataset, TS_synth_, was generated following the same procedure as DS_synth_ (see section 4.1) with 2000 images, but with enhanced saliency. Nuclei masks are provided by the input generation procedure (elliptical shapes).

Taking these 3 datasets together (summarized in table 1), the target style dataset contains 3119 images.

**Table 1:** Number of images and nuclei in each dataset (DS column) used for the training and testing of SalienceNet.

| DS | #Images | #Nuclei | Style |
| --- | --- | --- | --- |
| DS<sub>org</sub> | 1056 | 43633 | Source |
| DS<sub>synth</sub> | 1500 | 128962 | Source |
| TS<sub>LS</sub> | 568 | 20754 | Target |
| TS<sub>DB</sub> | 551 | 23121 | Target |
| TS<sub>synth</sub> | 2000 | 171915 | Target |

## 5 RESULTS

To train the SalienceNet models, we split each of the two source datasets as well as the two experimental target datasets (see figure 3) into train and test subsets, in 90% and 10% proportions. The test datasets are denoted 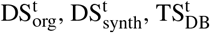 and 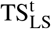, respectfully.

We then performed the hyperparameter search by exploring all possible combinations of α, β, and γ (weights of the loss components, see section 3.2) with step of 0.1 in order to estimate which combination of parameters yielded the best model. This resulted in 42 models, denoted by (α, β, γ) combinations in figure 4. Moreover, for comparison purposes we have trained a vanilla CycleGAN model, without any modification with respect to the original CycleGAN network.

**Figure 4:**
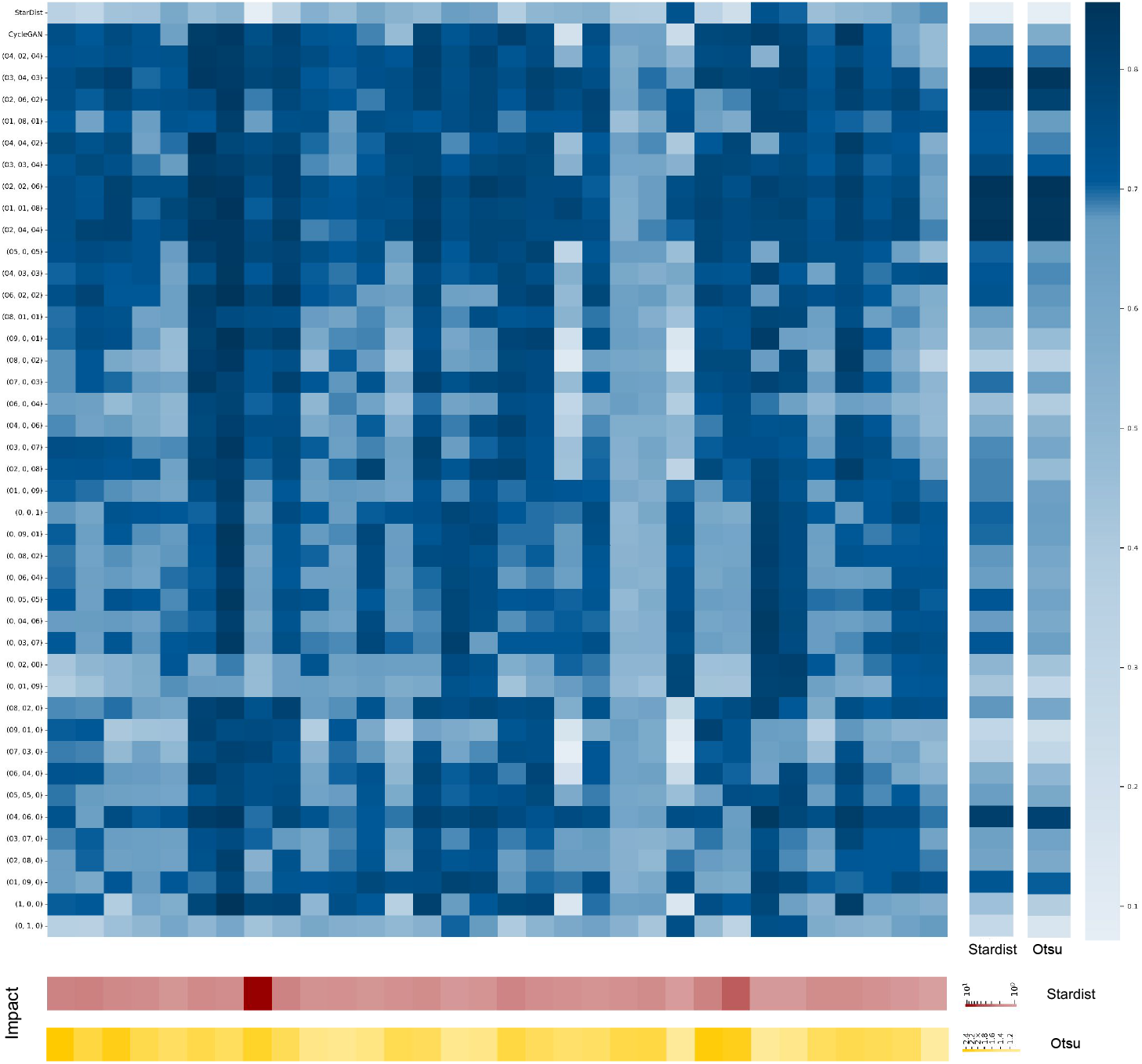
Heatmap of the IOU values for each image segmentation of 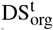, after enhancement and without enhancement. Average IOU values are shown in the two right columns (StarDist and Otsu). First line: IOUs obtained for non-enhanced images; second line: IOUs after enhancement by the vanilla CycleGAN and segmentation by StarDist, all the other lines: IOUs for segmentation by StarDist after enhancement by SalienceNet for different α, β and γ values. Bottom-most rows (red and yellow color-scale) show the impact of *M* enhancement on segmentation quality as IOU ratio (log scale).

All the 42 SalienceNet models and the vanilla CycleGAN were applied to the 4 test datasets 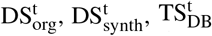 and 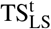 to perform saliency enhancement. The original images and their enhanced counterparts were then segmented, without any parameter tuning, using two widely used segmentation methods:

1. the non-parametric version of the classical segmentation Otsu thresholding method with an adaptive threshold,
2. StarDist, a deep-learning based segmentation, with the 2D fluo versatile model as provided by (Schmidt et al., 2018), without re-training on our data or supplementary fine-tuning.

The resulting masks were then compared with ground truth. For this purpose, expert ground truth annotation was performed on the 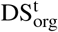 test dataset; nuclei masks (ground truth) were already available for the 3 other test datasets (see sections 4.1 and 4.2).

To measure the quality of the resulting segmentation, we computed the intersection over union (IOU) for each image to quantify the overlap (in pixel count) between the segmentation and the ground truth: 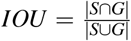, where *S* is the mask resulting from segmentation and *G* is the ground truth mask.

### SalienceNet enables accurate nuclei segmentation

First, we determined the most performant model with respect to our goal of segmenting low SNR images from live-cell imaging by looking at the enhancement performance on organoid images. Figure 4 shows the IOU scores for segmentation by StarDist of both enhanced (by 42 models) and non-enhanced images, and that for each image of the experimental organoid test dataset 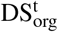. Individual model results for Otsu segmentation being very similar (albeit slightly worse), are not shown in this figure.

The IOU is shown in figure 4 for each image with respect to the ground truth. The geometric mean of all nonenhanced image’s IOU was 0.49 for StarDist segmentation and 0.45 for Otsu segmentation, while the geometric mean for IOU after saliency enhancement ranged from 0.54 to 0.75 for StarDist and from 0.48 to 0.75 for Otsu segmentation. Best results were obtained for segmentation after enhancement by the SalienceNet model with α = 0.2, β = 0.2, and γ = 0.6, with a geometric mean of IOU of 0.75 for both StarDist and Otsu. We denote this model by *M*.

The impact of this model *M* on the quality of the downstream segmentation was computed as the ratio of IOU for images enhanced with *M* over the IOU of non-enhanced images. Impact values range between 1.08 and 2.57 for Otsu segmentation and between 1.09 and 10.73 for StarDist segmentation; notice that the lower bound is *>* 1 in both cases.

An illustration of StarDist segmentation results for non-enhanced images and for those enhanced by *M* for low saliency organoid images is provided in figure 5.

**Figure 5:**
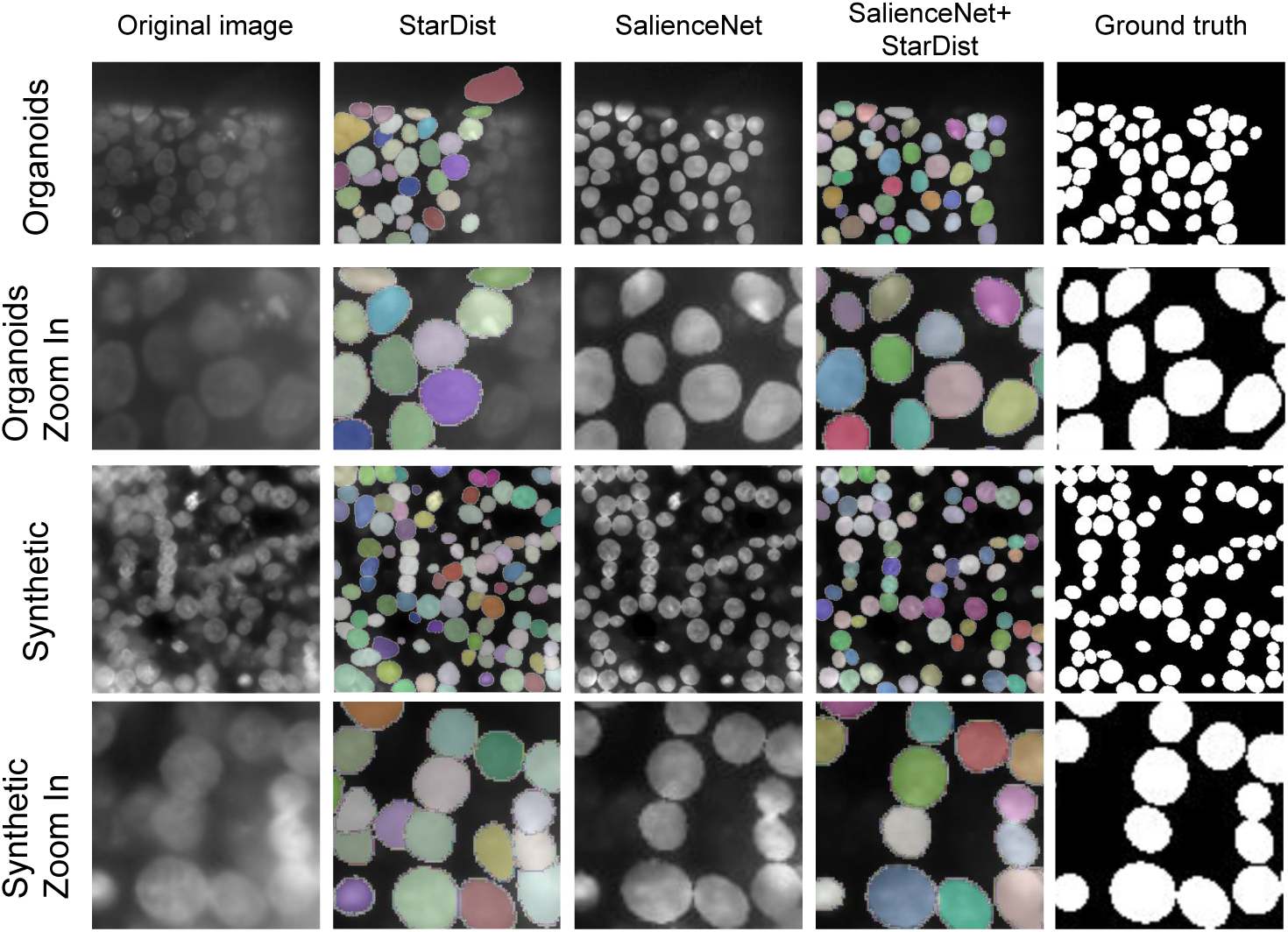
Examples of segmentation results by StarDist obtained without enhancement and after enhancement by SalienceNet *M* model. Sample images come from two low saliency test datasets: 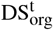 for the two upper rows, and 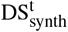 for the two lower rows. Rows 2 and 4 show a zoom-in of the respective rows right above.

Table 2 shows the IOU values for the best model *M* of SalienceNet applied to the 4 test datasets before and after enhancement by SalienceNet. On one hand, we observed that SalienceNet indeed improved the accuracy of nuclei segmentation in low-light source datasets. On the other hand, this table shows that for the already salient images, SalienceNet did not degrade the quality of segmentation.

**Table 2:** IOU values for segmentation with Otsu or StarDist of non-enhanced and enhanced with *M* SalienceNet model images in 4 test datasets. All reported values are the geometric mean, per dataset, of the IOU of individual images.

| **IOU**<br>**DS** | **Otsu** | **SN + Otsu** | **StarDist** | **SN + StarDist** |
| --- | --- | --- | --- | --- |
| $DS_{org}^t$ | 0.45 | 0.75 | 0.49 | 0.75 |
| $DS_{synth}^t$ | 0.62 | 0.90 | 0.62 | 0.86 |
| $TS_{LS}^t$ | 0.82 | 0.86 | 0.90 | 0.90 |
| $TS_{DB}^t$ | 0.69 | 0.78 | 0.83 | 0.83 |

Together, these results show that saliency enhancement by SalienceNet enables accurate downstream nuclei segmentation by widely used methods without need for parameter tuning.

### SalienceNet enhances nuclei saliency

To evaluate whether SalienceNet improved saliency, we computed its indirect measure - the Signal to Noise Ratio (SNR) as *SNR* = (*m − µ*_*B*_)*/*σ, where *m* is the maximum pixel intensity within the nuclei masks in an image, *µ*_*B*_ is the mean value of the background and σ is the standard deviation of the background. The SNR was computed for the source style datasets DS_org_ and DS_synth_, and the target style datasets TS_synth_, TS_LS_ and TS_DB_. We also measured the SNR of the test datasets 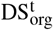 and 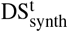 after enhancement by SalienceNet.

We observed (see figure 6) that SalienceNet enhanced SNR in low-light images of 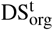 and 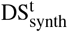 close to the level of SNR of the already salient experimental target style images TS_LS_ and TS_DB_, and up to the SNR level of the synthetic salient dataset TS_synth_.

**Figure 6:**
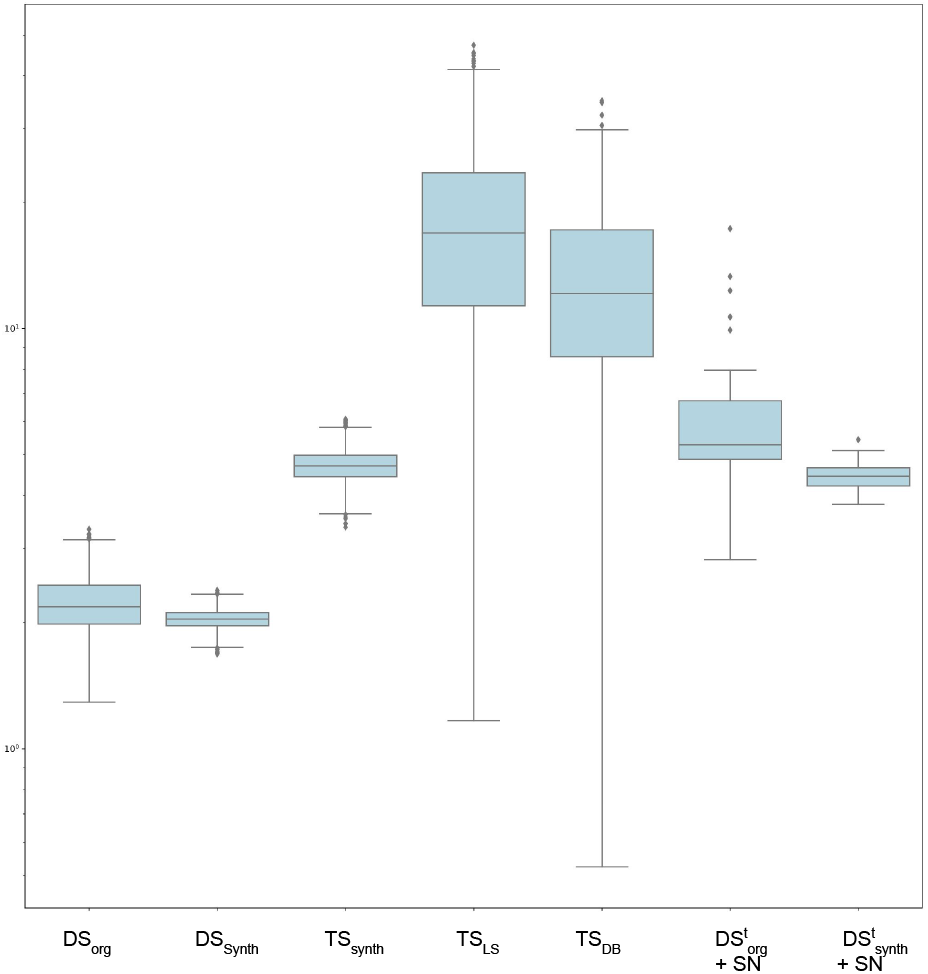
Signal to Noise Ratio (SNR) distributions. Boxplots represent the distribution of SNR for the source style images 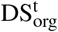 and 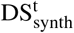 and the target style images TS_LS_ TS_DB_, TS_synth_. SNR distributions of the test images after enhancement with SalienceNet *M* model are shown for the low-light 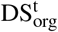 and 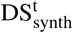 datasets.

## 6 SUMMARY

In this work, we introduced SalienceNet, a CycleGAN-based network specifically designed to enhance nuclei’s saliency in low SNR images that does not require annotation for training on new data.

We used the soSPIM light-sheet microscopy, a technique that allows to illuminate the biological sample with little light compared to other methods, to acquire organoid images. The result is that the illumination and the SNR are lower in these images and the nuclei are less salient. We used these organoid images as source style for training our network and further for testing. To implement SalienceNet we combined three loss functions with different properties and have shown that our adaptation of CycleGan improved segmentation results after enhancement relative to both segmentation of non-enhanced images and of those enhanced with the vanilla CycleGan.

We compared the segmentation results of widely used non-parametric Otsu thresholding and StarDist on both raw images and images enhanced with SalienceNet of our novel organoid live-cell imaging dataset. We have shown that using SalienceNet improved the segmentation quality of both classical and deep learning based nuclei segmentation algorithms in low SNR nuclei images. It should be noted that adding the SalienceNet enhancement step prior to nuclei segmentation did not degrade the quality of results for the already salient datasets. Taken together, these results show that SalienceNet is a useful new step for nuclei segmentation workflows.

## CODE AVAILABILITY

SalienceNet network’s code for training and testing nuclei enhancement is fully open source and available on GitHub at https://github.com/cbib/SalienceNet. Our best pre-trained model *M* used in this study is also available from the same GitHub page.

